# Universal patterns of matter and energy fluxes in land and ocean ecosystems

**DOI:** 10.1101/2020.04.13.039156

**Authors:** A. V. Nefiodov

## Abstract

In work [1], the fundamental relationships for the fluxes of matter and energy in terrestrial ecosystems were obtained. Taking into account the universal characteristics of biota, these relationships permitted an estimate to be made of the vertical thickness of the live biomass layer for autotrophs and heterotrophs. The distribution of consumption of biota production as dependent on the body size of heterotrophs was also investigated. For large animals (vertebrates), the energy consumption in sustainable ecosystems was estimated to be of the order of one percent of primary production. In this comment, it is shown that the results of work [1] also hold true for ocean ecosystems and thus are universal for life as a whole. This is of paramount importance for human life on Earth.

## On the system of units and dimensions of physical quantities

Since the density of living organisms practically coincides with the density of liquid water *ρ* = 1 t/m^3^, live body mass *m* and characteristic body volume *l* ^3^ are related as *m* = *ρl* ^3^. The universality of density *ρ* allows one to reduce the number of dimensions. Assuming by definition that *ρ* ≡ 1, one has *m* = *l*^3^, so that 1 g of live mass = 0.1 gC = 1 cm^3^. The second equality is written, taking into account that live mass contains about 10% of carbon C.

Similarly, in meteorology and hydrology, the relationship *m* = *l*^3^ allows one to relate the mass of precipitation to its volume, so that the amount of precipitation falling on a unit area of the Earth’s surface per unit time (for example, per year) acquires the dimension of velocity (mm/yr):

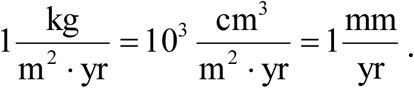

For biota, the energy content per unit of live mass is a universal characteristic [2]:

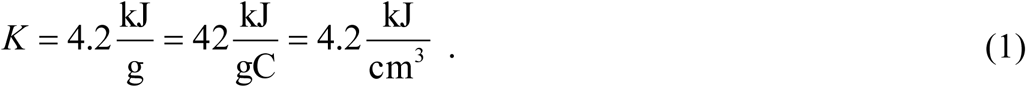

The quantity *K* makes it possible to measure the fluxes of organic substances and live mass in energy units. The notion of *surface-specific power of the energy flux* (or sometimes simply *specific power* with the dimension of W/m), as applied to living organism, means the amount of energy (J) transferred per unit time (s) through a unit area (m^2^) of the organism’s projection onto the Earth’s surface.

The decimal prefixes for fractions and multiples of units used in this work are: 10^−12^ pico (p), 10^−6^ micro (μ), 10^−3^ milli (m), 10^−2^ centi (c), 10^3^ kilo (k), 10^9^ giga (G).

## Energy fluxes in the ocean biota

According to the concept of biotic regulation [2,3], life on Earth exists sustainably by controlling the openness of nutrient cycles. The concentrations of organic and inorganic substances are kept by the biota in a stationary state optimal (most favorable) for life. In the absence of external environmental perturbations, the flux of synthesized organic matter is compensated by the reverse flux of inorganic substances, so that the nutrient cycles are closed. In the presence of external perturbations, living organisms function in such a way as to compensate for the emerging unfavorable changes in the concentrations of life-important substances and to return the environment to its original optimal state. It is achieved by directional deviations from the closeness of the biochemical cycles. The biosphere (global biota together with the external environment, surrounding it and interacting with it) maintains its sustainability only if the perturbations do not exceed a certain limit (threshold).

In work [1], within the framework of the concept of biotic regulation, several fundamental relationships for the fluxes of matter and energy in terrestrial ecosystems were studied. In this comment, these relationships are applied to oceanic ecosystems. The analysis is based on the laws of conservation of matter and energy, as well as on the existence of the optimal average mass-specific respiration rate of terrestrial and oceanic biota.

Oceanic phytoplankton mainly consists of single-celled organisms that synthesize the organic matter of their bodies from inorganic elements by absorbing solar photons (autotrophs). The reaction of photosynthesis can occur only in the uppermost (euphotic) layer of the ocean, where sunlight penetrates. The depth of this layer depends on the transparency of seawater, which is determined by the productivity of phytoplankton [4]. In the open ocean, this depth reaches an average of about 80 meters. Approximately 90% of solar radiation is already absorbed within the upper 20 meters. In contrast to the immotile vegetation cover of land, phytoplankton is randomly mixed by turbulent diffusion of the atmospheric wind power. The mixing is facilitated by the presence of a vertical temperature gradient (thermocline) and, accordingly, of a jump in the water density (pycnocline).

The organic matter of phytoplankton is consumed by heterotrophs that decompose it into inorganic substances. Oceanic heterotrophs include bacteria, as well as eukaryotic unicellular and multicellular organisms (zooplankton, i.e. passively floating animals). The larger representatives of zooplankton are able to swim actively in the vertical dimension. In addition, there is nekton (fish and marine mammals swimming in the 3-dimensional space of the ocean) and benthos (organisms inhabiting the ocean floor).

Life is organized in such a way that the decomposition of organic matter does not interfere with its synthesis. For this purpose, the direct and reverse processes in the ocean are spatially separated: the synthesis of organic substances is possible only in the euphotic layer, while their decomposition occurs throughout the entire depth of the ocean, including the seafloor sediments. In transparent water, the euphotic layer extends to a depth of *H* ∼ 200 meters and coincides with the epipelagic zone, where almost all live biomass of ocean is concentrated, including phytoplankton, zooplankton, and nekton.

The total (gross) primary production of photosynthesis in the ocean is divided between the respiration of phytoplankton and its net primary production, which is consumed by heterotrophs. A certain part of the net primary production is associated with the flux of nutrients from the oceanic depths to the epipelagic layer. It is referred to as new (or export) production.

Global values of net primary production *P*_*g*_ = 56 GtC/yr for land and *P*_*o*_ = 49 GtC/yr for the ocean are known from numerous independent measurements and estimates [5]. With the water surface area of *S*_*o*_ = 3.6·10^8^ km^2^, the net primary productivity of the ocean is given by

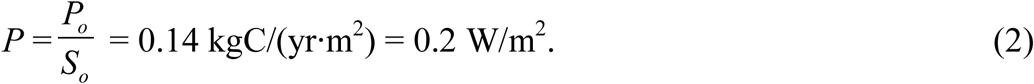

This value determines the average global power of the flux of organic synthesis per unit surface area. In the ocean, the value of *P* (2) is several times lower than on land, where it is estimated at about 0.5–1 W/m^2^ [1,4]. This difference is due to the strong absorption of the solar radiation in water [5]. However, since the surface area of the ocean is almost 2.5 times larger than the land area, the integral values of *P*_*o*_ and *P*_*g*_ approximately coincide, that is, the land and oceanic biota interact through a common atmosphere in a coherent manner, as ecological partners equal in power.

Across vast areas of the ocean, two genera of the cyanobacteria *Prochlorococcus* and *Synechococcus* with cell sizes of 0.5–0.7 μm and 0.8–1.5 μm, respectively, make the largest contribution to primary productivity [6]. Among heterotrophs, as a rule, unicellular eukaryotic zooplankton with a linear size of ∼ 1 to ∼ 200 μm and bacteria with a size of ∼ 0.3 to ∼ 1 μm dominate. Bacteria are the organisms decomposing the major part of the organic matter in the ocean. Importantly, these fluxes of matter pass through the detrital channel, that is, bacteria consume organic materials in the form of dead residues of other living organisms [7]. In tropical and subtropical regions, nutrients are mainly recycled in the epipelagic zone, while the contribution of new production is relatively small. Particles of phytodetritus and fecal pellets of microzooplankton sinking from the euphotic zone accumulate at the pycnocline (the level below which the seawater density increases stepwise), which plays the role of a “liquid seabed”.

More productive areas of the world’s oceans are located in coastal waters, in temperate and northern latitudes [8]. Here, in addition to bacteria with cell sizes of up to ∼ 1 μm, large phytoplankton with cell sizes of ∼ 5 to ∼ 100 μm and multicellular zooplankton with the body size of ≳ 200 μm are widespread. Particles of phytodetritus and fecal excretions of macrozooplankton can be quite massive to plunge deep into the ocean, passing through the pycnocline. The contribution of new production is significant due to the large flux of nutrients from the depths of the ocean (upwelling) and is comparable to the contribution from nutrient recycling within the euphotic layer.

According to the laws of matter and energy conservation, the power of photosynthesis *F* (W/m^2^) per unit of the oceanic surface area, in a stationary case, is spent on the respiration *J* (W/m^2^) of phytoplankton and its net primary productivity *P* (W/m^2^) [1]:

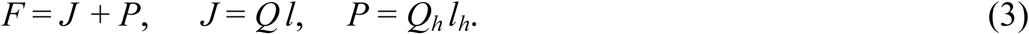

Here *J* (W/m^2^) *= Q* (W/m^3^) *· l*(m) denotes the respiration power, taken per unit of the horizontal surface area, of phytoplankton with its live biomass characterized by an effective horizontal layer of thickness *l*(m). Accordingly, *Q*_*h*_ and *l*_*h*_ are the respiration power per unit volume and the thickness of an effective horizontal layer of heterotrophs.

The respiration rates of the smallest phytoplankton and prokaryotic heterotrophs decomposing it are estimated as follows (see ref. [9], table 1):

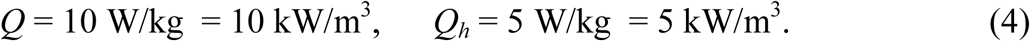

In a steady state, the phytoplankton layer of thickness *l = l*_*c*_ expends for respiration the power *J*_*c*_ = *Q l*_*c*_, which is approximately equal to the net primary productivity *P*_*c*_= *F* − *J*_*c*_ ≈ *J*_*c*_ [10,11]. Net primary productivity is completely consumed by the heterotroph layer of thickness *l*_*h*_:

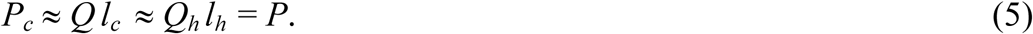

Accordingly, gross primary productivity can be written as

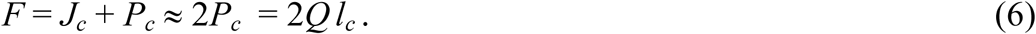

Taking into account estimate (2), it yields

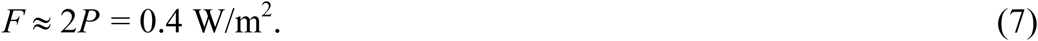

From formulas (4) and (5) one finds the thickness of the layers of phytoplankton and heterotrophs:

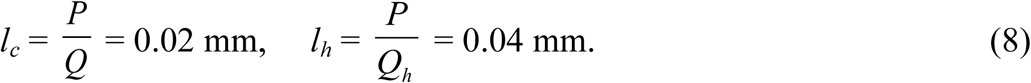

According to (5) and (6), the stationary energy flux of photosynthesis splits approximately equally between the metabolic powers (respiration) of autotrophs and heterotrophs, which are responsible for the oppositely directed fluxes of biological synthesis and decomposition of organic substances [1].

Taking into account Eqs. (1) and (5), time *τ = K/Q* of decomposition of the live mass of phytoplankton by respiration coincides with time *Kl*_*c*_ */P*_*c*_ of its synthesis in a layer of thickness *l*_***c***_:

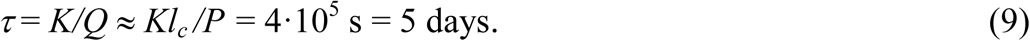

Due to the difference in respiration rates, the turnover time *τ*(9) of the biomass of phytoplankton is an order of magnitude shorter than that for the green leaves of plants on land [1].

Knowing the value of *l*_*c*_ (8) and the surface area *S*_*o*_ of the ocean, one obtains the volume of the live layer of phytoplankton *V*_*c*_ = *l*_*c*_*S*_*o*_ = 7 km^3^, which corresponds to the biomass of ∼ 0.7 GtC. This value is consistent within the margin of errors with the available modern experimental estimates for the biomass of cyanobacteria (∼ 0.3 GtC) and the total biomass of marine autotrophs (∼ 1.3 GtC) [12], most of which are microalgae. Similarly, from the layer thickness *l*_*h*_ (8) one can estimate the volume *V*_*h*_ = *l*_*h*_*S*_*o*_ = 14 km^3^ and biomass (∼ 1.4 GtC) of heterotrophs. According to the experimental data [12], the biomass of marine prokaryotic heterotrophs is estimated at ∼ 1.6 GtC, which contains the contributions from bacteria (∼ 1.3 GtC) and archaea (∼ 0.3 GtC). Note also that a large store (∼ 10 GtC) of virtually metabolically inactive benthic prokaryotes can play the function of a reserve biotic power, to become active during sudden environmental perturbations for their rapid compensation.

The respiration *Q* of the ocean’s smallest phytoplankton is an order of magnitude higher than the respiration of green leaves on land, however it is at the upper limit of the range of *optimal* values [9]. In addition, the specific power of the flux *P* (2) of organic synthesis in water is lower than on land. Due to these two factors, the thickness of the phytoplankton layer *l*_*c*_ (8) in the ocean turns out to be about 50 times less than the thickness of the live layer of terrestrial vegetation, which is equal to 1 mm [1]. In absolute terms, the live biomass of autotrophs in the ocean and green leaves on land differ by one order of magnitude and are estimated as ∼ 1.3 GtC and ∼ 15 GtC, respectively [12]. Since the layer *l*_*c*_ (8) should be distributed over the epipelagic zone to a depth of *H* ∼ 200 meters, the concentration of living phytoplankton becomes negligibly small. In addition, the short turnover time *τ* (9) of the biomass of unicellular phytoplankton coincides with the cell lifespan. In the result, unlike in the forests on land, there is no accumulation of living plant biomass in the ocean. This lack of food base makes the existence of large herbivorous animals in the open ocean impossible [1].

## Distribution of heterotrophs’ energy consumption by their sizes

Let us consider the distribution of energy consumption by the oceanic heterotrophs over the linear size *L* of their bodies (Fig. 1). The data on the global production of marine organisms with body masses *m* ranging from 10 μg to 1 t are taken from Table 1 of work [13]. The studied range of body mass was divided into two intervals: 10 μg ⩽ *m* ⩽ 1 g and 1 g ⩽ *m* ⩽ 1 t, corresponding to the intervals of body length 215 μm ⩽ *L* ⩽ 1 cm and 1 cm ⩽ *L* ⩽ 1 m. The power of consumption of organic matter by organisms from each interval was calculated by dividing the production of organisms from the interval by the efficiency *ε* of energy transfer from one trophic level to another, taken equal to *ε* = 0.125 as in work [13]. The total power of consumption of organic matter by all animals listed in Table 1 of work [13] turns out to be 8 GtC/yr, which corresponds to approximately 16.5% of the global value of net primary production *P*_*o*_ = 49 GtC/yr in the ocean [5]. This means that the main power of consumption of organic matter (∼ 83%, which corresponds to ∼ 41 GtC/yr) is accounted for by bacteria and microzooplankton with masses of 1 pg ⩽ *m* ⩽ 10 μg and body sizes of 1 μm ⩽ *L* ⩽ 215 μm. It is these smallest organisms that directly consume the major part of the net primary production synthesized by phytoplankton (see Fig. 1).

**Fig. 1.**
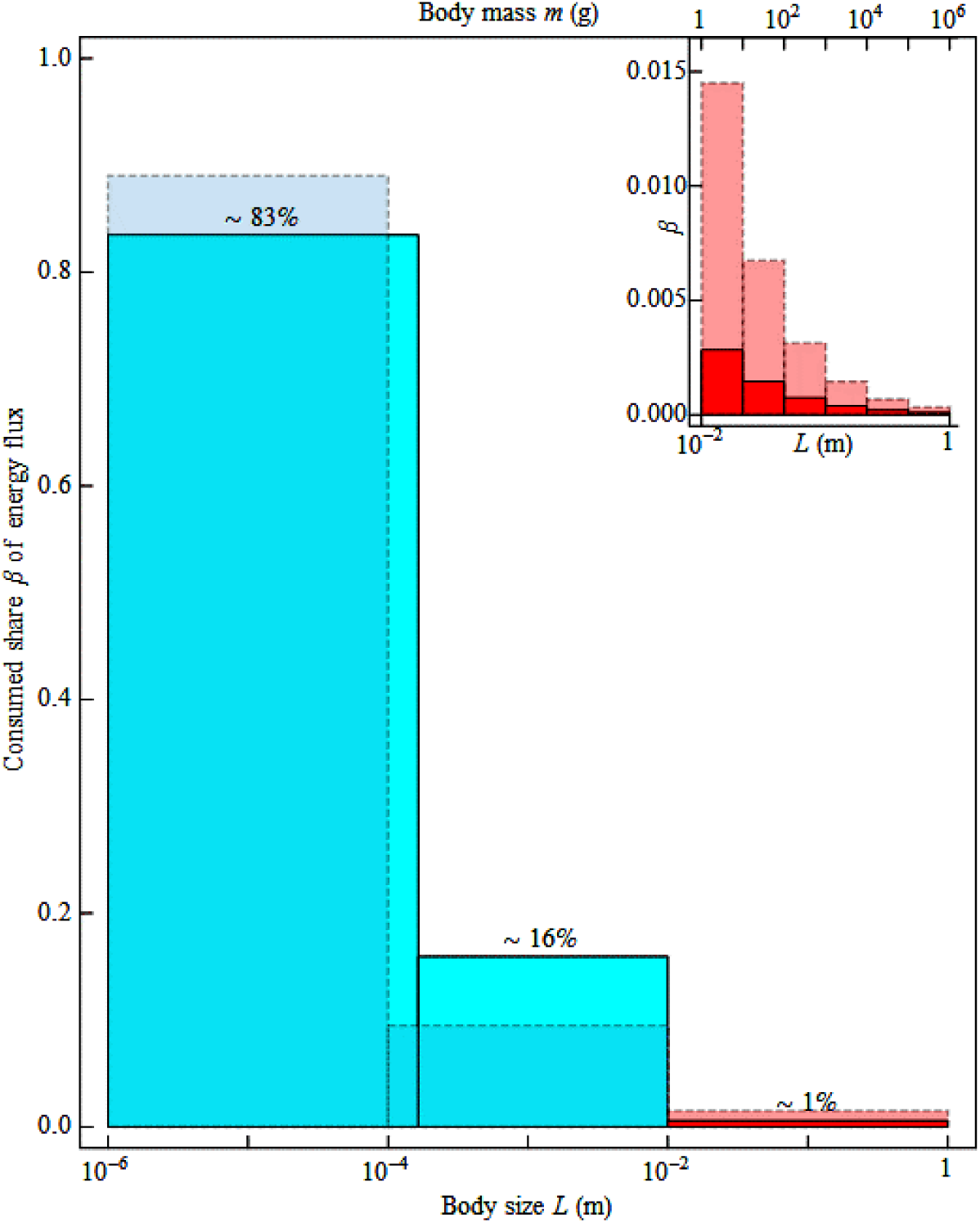
The share of consumption *β* of organic matter by heterotrophs in the ocean (solid lines, calculated in the present work) and in sustainable terrestrial ecosystems (dashed lines, taken from works [1,14,15]), depending on body size. The characteristic linear size *L* is related to live mass *m* through the relationship *L* (cm) = *m*^1/3^ (g^1/3^). The areas of hyperfine tuning corresponding to the largest body sizes are shaded in the hues of red.

The organisms with body sizes within the intervals of 215 μm ⩽ *L* ⩽ 1 cm and 1 cm ⩽ *L* ⩽ 1 m play the role of, respectively, fine and hyperfine tuning in ecosystem functioning, although the biomass of marine animals (∼ 2 GtC) [12] exceeds the total biomass of oceanic autotrophs. The latter circumstance is associated with the low metabolic activity of heterotrophs, due to the high cost of respiration in the viscid aqueous environment with a low content of oxygen [9].

In Fig. 1, the dashed line also shows for comparison the distribution of consumption of net primary production in terrestrial ecosystems undisturbed by anthropogenic impact: ∼ 90% is contributed by bacteria and saprophagous fungi, ∼ 10% by invertebrates, and ∼ 1% by vertebrates (primarily mammals) [1,14,15]. These two histograms have a similar structure and describe the universal pattern of energy fluxes in the sustainable natural biota. The discrepancy between the distributions for large organisms on land and in the ocean (graphs in the top right hand corner of Fig. 1) is likely caused by over-fishing.

For multicellular phytoplankton, the turnover time *τ* (9) of biomass does not coincide with the cell lifespan. The plant biomass can accumulate up to concentrations sufficient for direct consumption by macroscopic zooplankton with a body size of *L* ≳ 200 μm (middle interval of histogram in Fig. 1). These organisms have a significant impact on the productivity of local ecosystems by regulating the fluxes of dead organic matter passing through the pycnocline (and, accordingly, the fluxes of new production). On land, the middle interval of histogram in Fig. 1 is represented by invertebrate organisms, in particular insects, which are admitted for pollination of many plant species and thus also have a significant effect on the net primary productivity and thus on the overall ecosystem functioning.

## Paradox of the plankton

The originally described in work [16] is a lack of understanding how a wide variety of competing species of phytoplankton sustainably coexist in aquatic ecosystems with seemingly limited resources. This is still a hot topic of research, but further studies raise rather new questions only (see, for example, [17-19]).

In fact, the paradox of the plankton consists in an incorrect formulation of the problem. Without taking into account that it is the natural biota that is the main regulator of the environment, it is neither possible to understand these problems in laboratory conditions, nor to describe them in mathematical models. Only within the framework of the biotic regulation concept, the seemingly unconnected paradoxical phenomena can be combined into a single self-consistent picture. The variety of species is necessary to cope with the greatest challenge of maintaining the environment in a steady state. By virtue of the law of large numbers, the biota effectively minimizes the fluctuations of parameters of the environment, which is maintained by the biota in a highly non-equlibrium physico-chemical state. The environment is composed of “apparently limited resources” precisely because it is the most favorable for life. In such an optimal state, all species in the ecological community, through which the major part of the energy fluxes passes, respire at an optimal rate.

Small populations can only exist under very stable (weakly fluctuating) external conditions, since the probability of species survival declines with a decreasing number of individuals. In this case, the stability of external conditions is maintained by the biota itself due to the fast compensation of any below-threshold perturbations. In a stable ecosystem at fixed primary flux of solar radiation, more species with fewer individuals may exist (see [2], Section 3.9), which explains the rich species diversity of oceanic phytoplankton.

Next, the genetic program of the biotic regulation must be rigorously stabilized by natural selection [1]. This stabilization explains why, despite the enormous population numbers of some ubiquitous species of phytoplankton, such as *Emiliania huxleyi* (Haptophyta), these species have very low genetic variability (the so-called Lewontin’s paradox) [19].

## Conclusion

Born in ocean, life has not stopped since its origin. For example, according to the paleontological data, the earliest ecosystems appeared about 3.5 billion years ago were microbial mats (symbiotic communities of microorganisms, mainly consisting of prokaryotes), which in modern forms still inhabit the Earth [20]. The environment formed by the activity of prokaryotes [21].

To date, there is no evidence that biota has ever lost its sustainability and, accordingly, the control over its environment. Despite the mass extinctions of higher taxa of flora and fauna, which took place in the history of life [22], the environment remained suitable for life at all times. The key problem is not the changes in biodiversity as a result of natural disasters, but the unknown temporal dynamics of the productivity of global biota.

Currently, the gross power of the Earth’s biota is ∼ 200 GtC/yr [5]. In a steady state, it is spent on respiration of autotrophs and heterotrophs in approximately equal proportions. Unicellular organisms of microscopic dimensions channel up to ∼ 90% of the power of nutrient and energy fluxes through themselves and perform the main work of maintaining the environmental state optimal for life. Due to a huge number of living organisms [2,23], fluctuations in the fluxes of synthesis and decomposition are minimized relative to the value of dynamic equilibrium in accordance with the law of large numbers. Note that the first estimate of the number of cells in the biosphere was obtained by V. G. Gorshkov [2].

Man by his ecologically permitted share of the consumption of biosphere production is similar to a large vertebrate animal. As such, man is limited to the area of hyperfine tuning (Fig. 1). Since the global power of biota available to heterotrophs is ∼ 100 GtC/yr [5], the consumption of the total matter and energy fluxes at the threshold level of ∼ 1% is estimated as ∼ 1 GtC/yr. This includes human food, livestock fodder, and forest wood consumption. Currently, the maximum permissible limit is exceeded by an order of magnitude [2,14,24], largely due to the high population numbers and the destruction of natural forests.

This violation in the distribution of energy consumption in terrestrial ecosystems has led to the violation of the observed environmental sustainability and to the degradation of climate. Phytoplankton has such a low concentration of live biomass that humanity has not been able to exploit it yet. Therefore, the basis of energetics of ocean ecosystems has remained relatively undisturbed. The power of biotic regulation today is thus provided by ocean ecosystems and intact land ecosystems, mainly natural forests.

In philosophy, there is a statement “*freedom is the appreciation of necessity”* that is attributed to many authors (see, for example, [25], page 140) and interpreted in different ways. In the biotic regulation context, the existence of a fundamental restriction on the consumed fluxes of matter and energy means the primacy of the laws of nature, which humanity should harmonize its needs with. Only by restricting consumption below the threshold level and, accordingly, by reducing the current population numbers, will humanity be able to live and develop freely on our planet unbothered by environmental problems. The sustainability of the optimal environment and climate will be provided by the natural biota of the Earth, because it is the only power capable of solving that task.

## References

1. Gorshkov V. G., Makarieva A. M. Key ecological parameters of immotile versus locomotive life. Russian Journal of Ecosystem Ecology. 2020, vol. 5 (1), pp. 1–18. DOI: 10.21685/2500-0578-2020-1-1.

2. Gorshkov V. G. Physical and biological bases of life stability. Man, biota, environment. Berlin: Springer, 1995, 340 p. DOI: 10.1007/978-3-642-85001-1.

3. Gorshkov V. G., Gorshkov V. V., Makarieva A. M. Biotic regulation of the environment: Key issue of global change. London: Springer, 2000, 368 p.

4. Whittaker R. H. Communities and Ecosystems. New York: MacMillan Publishing Co. Inc., 2nd edition, 1975, 385 p.

5. Field C. B., Behrenfeld M. J., Randerson J. T., Falkowski P. Primary Production of the Biosphere: Integrating Terrestrial and Oceanic Components. Science. 1998, vol. 281 (5374), pp. 237–240. DOI: 10.1126/science.281.5374.237.

6. Flombaum P., Gallegos J. L., Gordillo R. A., Rincón J., Zabala L. L., Jiao N., Karl D. M., Li W. K. W., Lomas M. W., Veneziano D., Vera C. S., Vrugt J. A., Martiny A. C. Present and future global distributions of the marine Cyanobacteria *Prochlorococcus* and *Synechococcus*. Proceedings of the National Academy of Sciences U.S.A. 2013, vol. 110 (24), pp. 9824–9829. DOI: 10.1073/pnas.1307701110.

7. Toggweiler J. R. Diving into the organic soup. Nature. 1990, vol. 345, pp. 203–204. DOI: 10.1038/345203a0.

8. Huston M. A., Wolverton S. The global distribution of net primary production: resolving the paradox. Ecological Monographs. 2009, vol. 79 (3), pp. 343–377. DOI: 10.1890/08-0588.1.

9. Makarieva A. M., Gorshkov V. G., Li B.-L., Chown S. L., Reich P. B., Gavrilov V. M. Mean mass-specific metabolic rates are strikingly similar across life’s major domains: Evidence for life’s metabolic optimum. Proceedings of the National Academy of Sciences U.S.A. 2008, vol. 105 (44), pp. 16994–16999. DOI: 10.1073/pnas.0802148105.

10. Whittaker R. H., Likens G. E. The Biosphere and Man. In Series: Lieth H., Whittaker R. H. (Eds.) Primary Productivity of the Biosphere. Ecological Studies (Analysis and Synthesis), vol. 14. Berlin: Springer, 1975, pp. 305–328. DOI: 10.1007/978-3-642-80913-2_15.

11. Arístegui J., Harrison W. G. Decoupling of primary production and community respiration in the ocean: implications for regional carbon studies. Aquatic Microbial Ecology. 2002, vol. 29, pp. 199–209. DOI: 10.3354/ame029199.

12. Bar-On Y. M., Phillips R., Milo R. The biomass distribution on Earth. Proceedings of the National Academy of Sciences U.S.A. 2018, vol. 115 (25), pp. 6506–6511. DOI: 10.1073/pnas.1711842115.

13. Jennings S., Mélin F., Blanchard J. L., Forster R. M., Dulvy N. K., Wilson R. W. Global-scale predictions of community and ecosystem properties from simple ecological theory. Proceedings of the Royal Society B: Biological Sciences. 2008, vol. 275 (1641), pp. 1375–1383. DOI: 10.1098/rspb.2008.0192.

14. Gorshkov V. G. The Distribution of Energy Flows among the Organisms of Different Dimensions. Journal of General Biology. 1981, vol. 42 (3), pp. 417–429. [In Russian]

15. Makarieva A. M., Gorshkov V. G., Li B.-L. Body size, energy consumption and allometric scaling: A new dimension in the diversity-stability debate. Ecological Complexity. 2004, vol. 1 (2), pp. 139–175. DOI: 10.1016/j.ecocom.2004.02.003.

16. Hutchinson G.E. The Paradox of the Plankton. The American Naturalist. 1961, vol. 95 (882), pp. 137–145. http://www.jstor.org/stable/2458386

17. Roy S., Chattopadhyay J. Towards a resolution of =*the paradox of the plankton*’: A brief overview of the proposed mechanisms. Ecological Complexity. 2007, vol. 4 (1-2), pp. 26–33. DOI: 10.1016/j.ecocom.2007.02.016.

18. Cuesta J. A., Delius G. W., Law R. Sheldon spectrum and the plankton paradox: two sides of the same coin—a trait-based plankton size-spectrum model. Journal of Mathematical Biology. 2018, vol. 76, pp. 67–96. DOI: 10.1007/s00285-017-1132-7.

19. Filatov D. A. Extreme Lewontin’s Paradox in Ubiquitous Marine Phytoplankton Species. Molecular Biology and Evolution. 2019, vol. 36 (1), pp. 4–14. DOI: 10.1093/molbev/msy195.

20. Microbial Mats: Modern and Ancient Microorganisms in Stratified Systems. In Series: Seckbach J., Oren A. (Eds.) Cellular Origin, Life in Extreme Habitats and Astrobiology, vol. 14. Springer Netherlands, 2010, 606 p. DOI: 10.1007/978-90-481-3799-2.

21. Katola V. M. Ancient Procariotes: Origin, Evolutionary Path and Role in Earth’s History (Review). Bulletin Physiology and Pathology of Respiration. 2014, (52), pp. 129–135. [In Russian]

22. Raup D. M., Sepkoski J. J. Jr. Mass Extinctions in the Marine Fossil Record. Science. 1982, vol. 215 (4539), pp. 1501–1503. DOI: 10.1126/science.215.4539.1501.

23. Whitman W. B., Coleman D. C., Wiebe W. J. Prokaryotes: The unseen majority. Proceedings of the National Academy of Sciences U.S.A. 1998, vol. 95 (12), pp. 6578–6583. DOI: 10.1073/pnas.95.12.6578.

24. Vitousek P. M., Ehrlich P. R., Ehrlich A. H., Matson P. A. Human Appropriation of the Products of Photosynthesis. BioScience. 1986, vol. 36 (6), pp. 368–373. DOI: 10.2307/1310258.

25. Engels F. Herr Eugen Dühring’s Revolution in Science (Anti-Dühring). Moscow: Progress Publishers, 1977, 520 p.

